# Piezo1 is a mechanosensor of soft matrix viscoelasticity

**DOI:** 10.1101/2024.06.25.600570

**Authors:** Mariana A. G. Oliva, Giuseppe Ciccone, Jiajun Luo, Jonah L. Voigt, Patrizia Romani, Oana Dobre, Sirio Dupont, Massimo Vassalli, Manuel Salmeron-Sanchez

**Affiliations:** Centre for the Cellular Microenvironment (CeMi), University of Glasgow, The Advanced Research Centre, Glasgow, UK; Department of Molecular Medicine (DMM), University of Padua, Padua, Italy; Institute for Bioengineering of Catalonia (IBEC), The Barcelona Institute for Science and Technology (BIST), 08028 Barcelona, Spain; Institució Catalana de Recerca i Estudis Avançats (ICREA), Barcelona, Spain; Max Planck Institute for Medical Research, Heidelberg, Germany; Cellular Biomechanics, Faculty of Engineering, Bayreuth University, Bayreuth, Germany

## Abstract

Mechanosensitive ion channels have emerged as fundamental proteins in sensing extracellular matrix (ECM) mechanics. Among those, Piezo1 has been proposed as a key mechanosensor in cells. However, whether and how Piezo1 senses time-dependent ECM mechanical properties (i.e., viscoelasticity) remains unknown. To address this question, we combined an immortalised mesenchymal stem cell (MSC) line with adjustable Piezo1 expression with soft (400 Pa) and stiff (25 kPa) viscoelastic hydrogels with independently tuneable Young’s modulus and stress relaxation. We demonstrate that Piezo1 is a mechanosensor of viscoelasticity in soft ECMs, consistent with the molecular clutch model. By performing RNA sequencing (RNA-seq), we identified the transcriptomic phenotype of MSCs response to matrix viscoelasticity and Piezo1 activity, highlighting gene signatures that drive MSCs mechanobiology in soft and stiff viscoelastic hydrogels.

## Introduction

It is now widely accepted that the mechanical properties of the ECM drive cellular behaviour such as differentiation, proliferation, and migration^1–3^. The dynamic molecular interactions between the cell and its surrounding matrix have been explained by the existence of a molecular clutch between the cell’s actin cytoskeleton, myosin contractile motors and the adaptor proteins that connect ECM binding integrins to the actin cytoskeleton. The model was first introduced by Chan and Odde^4^, and since then, cell response to matrix stiffness (also referred to as rigidity)^5,6^, viscosity^7^ and viscoelasticity^8,9^ have been described via these mechanisms in more evolved versions of the framework.

Studies of cell response to different matrix mechanics therefore focus on mechanisms of cell-ECM adhesion, adaptive structural proteins, and molecules of the ECM^10^. Of note, transmembrane ion channels have been highlighted for their role in mechanosensing_11_. Specifically, the Piezo1 channel has quickly become a point of reference for mechanobiology studies. In 2010, the channel was identified in a high-throughput screening for integrin co- activators in epithelial cells and later categorised as a mechanically activated cation channel by Ardem Patapoutian and colleagues^12,13^. Recently, a concerted relationship between Piezo1 and integrin-mediated focal adhesion (FA) signalling has been described^14–16^, which has proposed Piezo1 channel activity as a key mediator of integrin signalling, and therefore of cell-matrix interaction.

Previous studies have found that substrate stiffness alone was sufficient to enhance Piezo1 mediated Ca^2+^ signalling, which in turn influenced the differentiation of neural stem cells^17^. Substrate elasticity has shown to drive cell function via the engagement of the actin-talin-integrin-fibronectin molecular clutch^6^, suggesting that mechanotransduction is dependent upon a stiffness threshold which promotes clutch engagement. Because of this, and the coordinated action of Piezo1 activity and integrin signalling^18^, we hypothesised that Piezo1 expression would affect molecular clutch engagement and downstream mechanotransduction in mesenchymal stem cells (MSCs).

However, native ECMs do not behave as perfectly elastic solids, instead, when assessing their response to mechanical deformations, reconstituted ECMs initially resist deformation followed by a time-dependent energy dissipation, characteristic of viscoelastic materials^8,18,1^. Energy dissipation arises from the dynamic molecular organisation of the ECM, which is not a perfectly chemically crosslinked network, but instead showcases breaking of weak bonds^19^, protein unfolding^20^ and entanglement release^8,21^. The interaction of cells with a viscoelastic substrate has been modelled via the same molecular clutch mechanism, however how and whether Piezo1 senses ECM viscoelasticity within this framework remains unknown.

In our work, we used two pairs of viscoelastic polyacrylamide (PAAm) hydrogels^22,23^ with either a low or high elastic component (Young’s modulus *E* = 400Pa and 25kPa)^27^, each with a higher or lower dissipative component, respectively. By combining these hydrogels with a mechanosensitive immortalised stem cell line in which we could tune Piezo1 expression, we demonstrated that Piezo1 mediates viscoelasticity sensing within the molecular clutch framework, especially at low substrate stiffness. Specifically, enhanced energy dissipation at low stiffness promotes cell spreading, focal adhesion formation, and overall molecular clutch engagement in a Piezo1 dependent manner. These results were consistent with downstream mechanotransduction events including enhanced cell metabolic capacity and transcriptional adaptation. Using RNA sequencing (RNAseq), we identify differentially regulated genes mediating cell response to substrates stiffness, viscoelasticity, and Piezo1 expression. Our results identify Piezo1 as a mechanosensor of time-dependent ECM mechanics in addition to substrate stiffness.

## Results and Discussion

### Cell response to matrix viscoelasticity is Piezo1-dependent

Two different hydrogel pairs were synthesised via mixing different amounts of acrylamide (AAm) and bis-acrylamide (BisAam) to obtain hydrogels of approximately the same *E* but varying stress relaxation rate. To achieve soft (*E* ≈ 400 Pa) and stiff (*E* ≈ 25 kPa) hydrogels, we combined two previously reported strategies to tune viscoelasticity in PAAm hydrogels (**Fig. 1, a**). For the stiff pair, the method first reported by Cameron and colleagues^26^ and later optimised by our group^28^ was employed. Here, substrate viscoelasticity is mediated by the movement of loosely crosslinked polymer chains. To create a softer pair of viscoelastic gels, the approach reported by Charrier and colleagues^23,24^ was adopted, as the first strategy gave rise to sticky hydrogels that were difficult to handle. In this case, substrate viscoelasticity arises from the physically entrapped chains of high molecular weight linear AAm. The resulting hydrogel groups presented a Young’s modulus of approximately 400 Pa and 25 kPa (hereafter referred to as *soft* and *stiff,* respectively); with no significant differences in Young’s modulus within each stiffness group (**Fig. 1, b**). To characterise the differences in the hydrogel’s stress relaxation behaviour, stress relaxation measurements were performed with a physiologically relevant step strain (*ε*) of 7%^24,25^ applied over 60 seconds using nanoindentation. From the resulting stress relaxation curves (**Fig. 1, c, d**) the time for the stress to relax to 80% of the initial value was calculated (**Fig. 1, e**), as well as the % of energy dissipation for each hydrogel condition (**Supplementary Data Fig. 1 c**). Resulting data demonstrated that for each stiffness group, there was a slow-relaxing (V-, elastic) and fast-relaxing (V+, viscoelastic) hydrogel, where the V+ hydrogels relax ∼ 2 times faster than their elastic counterparts (**Fig. 1, c**), as well as display higher relaxation amplitude (**Supplementary Data Fig. 1, c)**. This allowed for the investigation of cell response to substrate viscoelasticity independently of substrate elasticity, or Young’s modulus, in two distinct stiffness regimes in 2D. To confirm data obtained by nanoindentation, bulk rheology measurements were performed. By computing the ratio between the loss modulus (*G’’*) and the storage modulus (*G’*), we observed that the tan (δ) of the V+ hydrogels increased for both stiffness groups with respect to V- hydrogels (**Supplementary Data Fig. 1, d, e**), emulating soft tissues that exhibit loss moduli of approximately 10% of their elastic moduli at 1Hz^8^. Notably, as PAAm hydrogels are chemically crosslinked, both strategies give rise to viscoelastic solids with no plastic deformation, in contrast to physically crosslinked viscoelastic hydrogels^26^.

**Fig. 1.**
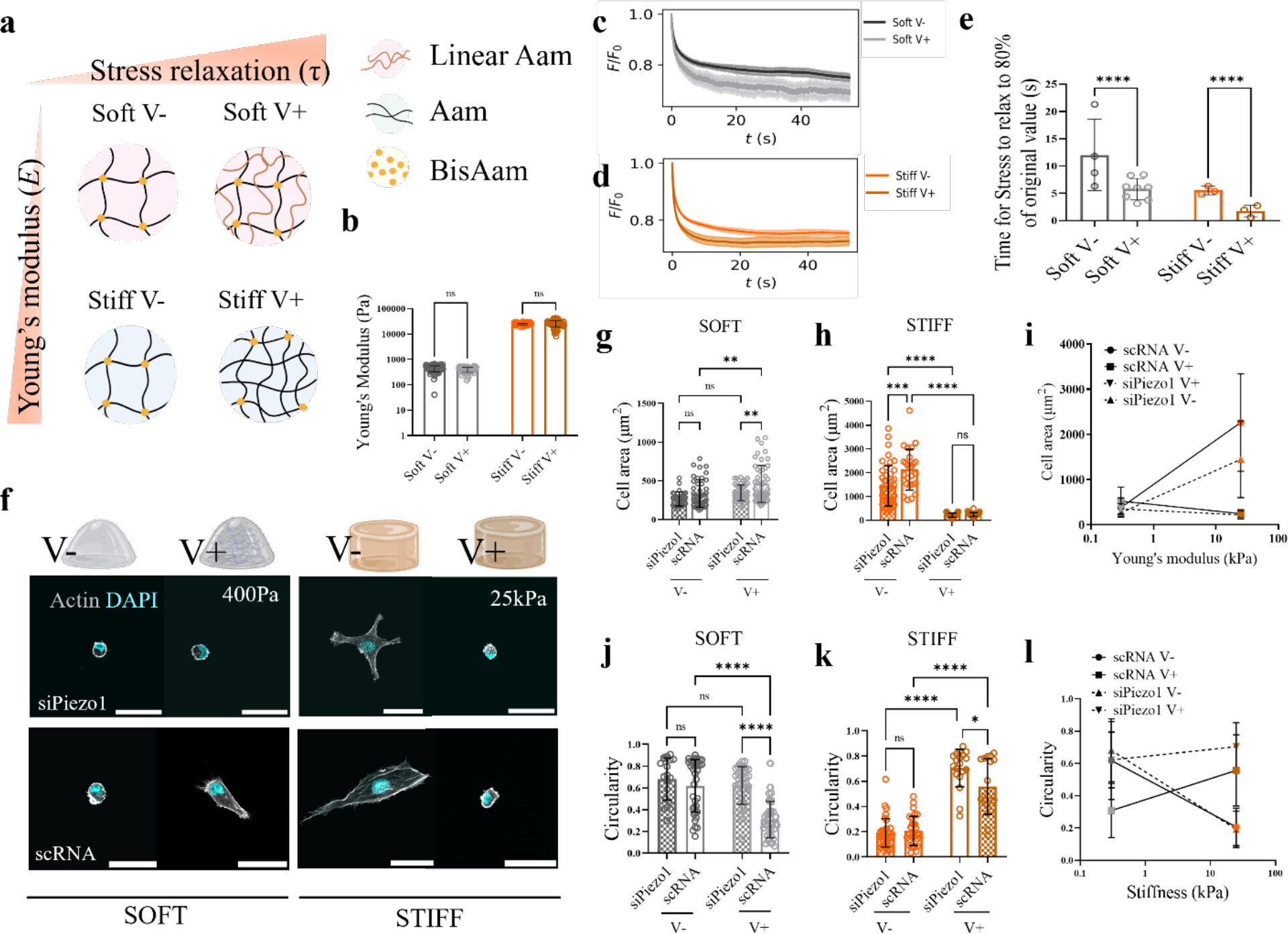
**Cell response to matrix viscoelasticity is Piezo1 dependent**. (**a**) Representation of hydrogel networks. (**b**) Nanoindentation measurements Young’s modulus data obtained for Soft (grey) and Stiff (orange) hydrogel pairs (n ≥ 51 single indentation curves) (**c**) Representative average stress relaxation curve ± SD of an indentation map performed on soft and stiff (**d**) hydrogels. (**e**) Time for the stress to relax to 80% of original value (s) plotted for Soft (grey) and Stiff (orange) hydrogel groups. (n ≥ 61 curves, each dot represents a map of ≥ 4 single nanoindentation curves each). (**f**) Representative immunofluorescence images of scRNA and siPiezo1 Y201 MSCs cultured on different hydrogel groups for 48h. Scale bar=50 µm (**g, h**) Quantified cellular area of the soft (left) and stiff (right) hydrogel groups. All individual points represent individual cell measurements. Data shown as mean ± SD (n ≥ 21). (**i**) Summary of mean cell area ± SD plotted as a function of Young’s modulus for all conditions. (**j-k**) Quantified circularity of the soft (left) and stiff (right) hydrogel groups. All individual points represent individual cell measurements. Data shown as mean ± SD (n ≥ 24). (**l**) Summary of mean circularity ± SD plotted as a function of stiffness for all conditions. Statistical analyses were performed using a one-way ANOVA (**b, c**) and two-way ANOVA test (**g, h, j and k**). P values indicating significance, ns > 0.05, * ≤ 0.05, ** ≤ 0.01, *** ≤ 0.001, **** ≤ 0.0001.

Next, we checked that the ECM matrix protein fibronectin (FN), the ECM element of the molecular clutch, would be homogeneous on all substrates regardless of stiffness and viscoelasticity, as previously reported^22,23^. By using the crosslinker sulfo-SANPAH, FN was conjugated on each hydrogel^27^. Through immunofluorescence, we confirmed a homogenous FN coating on all substrates with no significant changes in signal intensity (**Supplementary Data Fig. 1, a, b**). Overall, we have established a hydrogel system spanning a wide range of physiologically relevant stiffnesses with different rates of stress relaxation, allowing for the investigation of the effects of ECM viscoelasticity on cell behaviour independently of ligand density.

We then examined control (scRNA) and Piezo1 knock down (siPiezo1) Y201 MSCs (**Supplementary Data Fig. 2, a**) morphology on all hydrogel conditions (representative images in **Fig. 1, f**). Y201 MSCs showcased mechanosensitivity while also providing flexibility for the transient knock down of the Piezo1 channel with siRNA (**Supplementary Data, Fig. 2, b, c**). On soft hydrogels, scRNA cells increased their spreading area in response to substrate stress relaxation; whereas the opposite was observed for cells cultured on stiff substrates (**Fig. 1, g**). This was also reflected in terms of their circularity (**Fig. 1, j, k**). These data corroborate computational data first introduced by Chaudhuri and colleagues^9^, which suggested that cell spreading in response to increased substrate relaxation is stiffness-dependent and that would only increase at lower stiffness values in stress-relaxing substrates (<1pN/nm, or approximately 1kPa^28^). Piezo1 knock down has previously been reported to disrupt actin fibres and integrin activity, thus decreasing cell spreading^29^. This behaviour was also observed on Y201 MSCs cultured on fibronectin (FN) coated glass substrates (**Supplementary Data, Fig. 2, d, e and f**). Indeed, siPiezo1 cells demonstrated reduced cell spreading and increased circularity when compared to scRNA Y201 MSCs cultured on soft viscoelastic (V+) or stiff elastic (V-) hydrogels (**Fig. 1, f, j, k and l**). Interestingly, we also found that Piezo1 knock down abrogated cell spreading response to viscoelasticity at low stiffness regimes but not on stiff substrates, as cells presumably reached their minimum spreading area independently of Piezo1. Supporting this, cell spreading area was minimal on both soft elastic (V-) and stiff viscoelastic (V+) substrates (**Fig. 1, i**). Altogether, our results describe a Piezo1-dependent response to matrix viscoelasticity in soft matrices.

### Molecular clutch engagement in response to matrix viscoelasticity is Piezo1-dependent

To further investigate the mechanism of Piezo1 mediated ECM viscoelasticity sensing, we focused on quantifying actin-talin-integrin-fibronectin molecular clutch engagement. Clutch dynamics have been shown to regulate cell mechanotransduction in response to substrate stiffness^6^, viscosity^7^ and more recently, viscoelasticity^8,9^. However, how Piezo1 relays matrix viscoelastic cues via the molecular clutch is unknown. We first quantified focal adhesion (FA) formation by looking at vinculin through immunofluorescence, which recruits to adhesion sites in response to sustained force sensing by the actin-talin-integrin-fibronectin clutch^6^ (**Fig. 2, a**). Individual vinculin FA length (**Fig. 2 b, c**) and FA count (**Supplementary Data Fig. 3., b-d**) were quantified in soft and stiff groups. Data supported the previously observed cell spreading phenotype: faster substrate stress relaxation increases FA length in a soft regime (400Pa hydrogels), while the opposite was observed on stiff hydrogels (25kPa hydrogels) (**Fig. 2, d**).

**Fig. 2.**
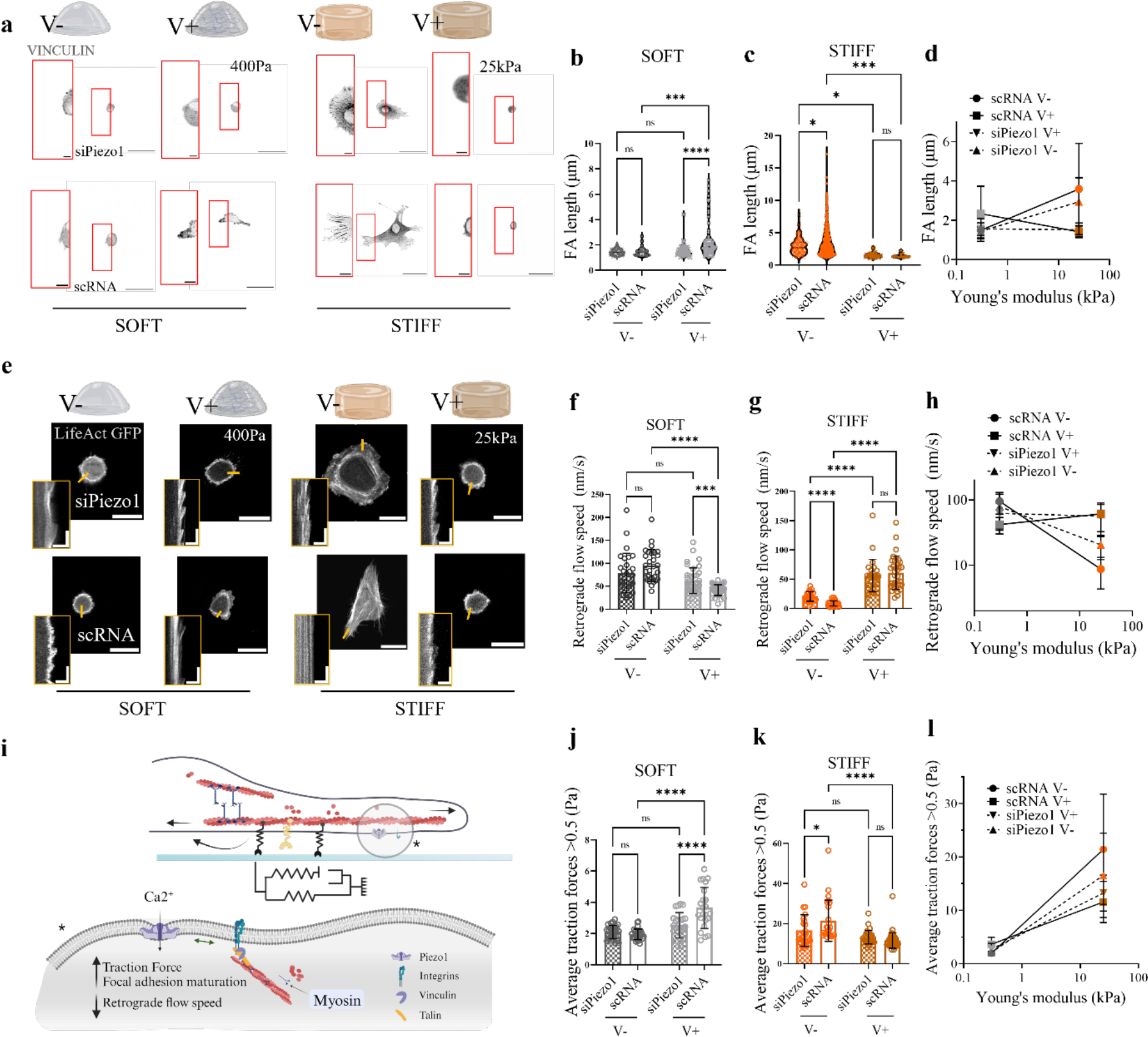
Molecular clutch engagement in response to matrix viscoelasticity is Piezo1-dependent. (**a**) Representative images of vinculin adhesions on siPiezo1 and scRNA Y201 cells on soft (left) and stiff (right) hydrogel groups of varying stress relaxation. Scale bar=50 µm, zoomed image scale bar=10 µm. (**b, c**) Quantified individual FA length of cells on the soft (left) and stiff (right) hydrogel groups. All individual points represent individual focal adhesion measurements. Data shown as mean ± SD (n ≥ 17). (**d**) Summary of mean individual FA length ± SD plotted as a function of Young’s modulus for all conditions. (**e**) Representative images of LifeAct GFP transfected siPiezo1 and scRNA Y201 cells on the soft (left) and stiff (right) hydrogel groups. Scale bar=50 µm. Insets are kymographs showing the movement of actin features, scale bar=5µm (horizontal) and 60s (vertical) (**f, g**) Quantified retrograde flow speed (nm/s) on the soft (left) and stiff (right) hydrogel groups. All individual points represent individual kymograph measurements. Data shown as mean ± SD (n ≥ 26). (**h**) Summary of mean retrograde actin flow ± SD plotted as a function of Young’s modulus for all conditions. (**i**) Diagram which describes the influence of the mechanosensitive channel Piezo1 in molecular clutch engagement. Created with BioRender.com (**j-k**) Average traction forces measured on siPiezo1 and scRNA Y201 cells cultured on the soft (left) and stiff (right) hydrogel groups (n ≥ 22). (**l**) Summary of average traction forces ± SD plotted as a function of Young’s modulus for all conditions. (**b-c, f-g, j-k**) Statistical analyses were performed using a two-way ANOVA test. P values indicating significance, ns > 0.05, * ≤ 0.05, *** ≤ 0.001, **** ≤ 0.0001.

Piezo1 activity has been intrinsically linked to FA dynamics since the channel’s identification in 2010 as an integrin co-activator^13–15,29^. Indeed, piezo1 knock down was sufficient to visibly reduce FA size and number in Y201 MSCs cultured on FN coated glass substrates (**Supplementary Data Fig. 2, e**). On soft fast-relaxing matrices (soft V+), siPiezo1 cells did not showcase increased adhesion size (**Fig. 2, b**) or number (**Supplementary Data Fig. 3, b**) when compared to the cells seeded on the elastic substrates (soft V-), highlighting the fundamental role of the channel in mediating cell response to matrix viscoelasticity at low substrate stiffness. In tandem, on stiff substrates, increased substrate relaxation (Stiff V+) significantly decreases FA length (**Fig. 2, c**) and number (**Supplementary Data Fig. 3, c**) on both siPiezo1 and scRNA Y201 MSCs. However, this effect is lessened on siPiezo1 cells, which showcase reduced adhesion structures when compared to the scRNA MSCs, as seen on cells cultured on glass substrates (**Supplementary Data Fig. 2, f**).

FA maturation and size are inversely related to the actin retrograde flow rate in cells^30^, as when the molecular clutch is engaged via actin-talin-integrin-fibronectin links, actin is bound in its fibrillar form and the rate of polymerisation decreases. Therefore, we transfected cells with Life-Act (**Fig. 2, e**) and measured the rate of actin retrograde flow using live confocal microscopy, in response to varying viscoelasticity on soft and stiff substrates and in terms of Piezo1 channel expression. In accordance with FA data, actin retrograde flow was slowed in response to faster substrate stress relaxation in soft hydrogels (**Fig. 2, f**), whereas on stiff hydrogels, actin retrograde flow increased ∼7 fold from a slow-relaxing (stiff V-) to a fast-relaxing (stiff V+) matrix in scRNA Y201 MSCs and ∼3 fold in siPiezo1 MSCs (**Fig. 2, g**). Notably, Piezo1 knock down abrogated any changes in retrograde flow speed between V- and V+ conditions in the soft group (**Fig. 2, h**). These data reiterate the role of Piezo1 in sensitively sensing the time-dependent mechanics of soft matrices and promoting clutch engagement in response to enhanced viscoelasticity on soft matrices.

To examine how the observed adhesion phenotype and actin polymerisation would translate in cell-exerted forces to the underlying matrix, we quantified the average cell traction forces exerted by scRNA and siPiezo1 cells on the substrates through traction force microscopy (TFM). Substrates were prepared including 200 nm fluorescent beads and the bead displacement before and after cell-applied force was calculated and converted into forces. We note that using an elastic algorithm likely overestimates forces on viscoelastic V+ substrates (**Supplementary Data Fig. 3, a**). Still, standard TFM has proven a useful tool to investigate relative changes in cell exerted forces on viscoelastic PAAm hydrogels^31^. Accordingly, we found that on soft substrates, increased substrate stress relaxation increased traction force generation in scRNA cells, whereas this increase was abrogated siPiezo1 Y201 MSCs (**Fig. 2**, **j**). On stiff substrates, both siPiezo1 and scRNA cells significantly decreased the average forces generated as substrate stress relaxation increased (**Fig. 2**, **k**). This response is lessened in siPiezo1 cells, which highlights the role of Piezo1 in traction force generation mechanisms within the cell, consistent with previous reports^16^. Perturbation studies also highlighted this relationship, as inhibiting cell contractility with blebbistatin demonstrated similar cell-substrate interaction phenotypes as those produced by siPiezo1 Y201 MSCs (**Supplementary Data Fig. 3, e-f**).

Overall, our results underscore the role of Piezo1 in mediating viscoelasticity-sensing in soft but not stiff ECMs, where channel knock down lessened cell response to increased substrate stress relaxation but did not fully revoke it. Indeed, our experimental data is in agreement with molecular clutch dynamics in response to surface viscoelasticity first described by Chaudhuri and colleagues^9^, where clutch engagement was enhanced by substrate stress relaxation on soft environments but inhibited above a stiffness threshold (∼1kPa)^9,28^. However, for the first time, we demonstrate how Piezo1 mediates clutch dynamics in cells interacting with a viscoelastic substrate (**Fig. 2, i**).

### Matrix viscoelasticity and Piezo1 regulate downstream mechanotransduction and mitochondrial morphology

We then investigated whether the observed changes in clutch engagement as a function of Piezo1 expression and substrate viscoelasticity would activate downstream mechanotransduction events. Increased cytoskeletal tension has been associated to nuclear compression and chromatin compaction^32^. Because of this, we first assessed and quantified nuclear projected spreading area (**Fig. 3**, **a-d**). On soft substrates (400 Pa), faster stress relaxation (Soft V+) promotes nuclear spreading compared to elastic hydrogels (Soft V-) (**Fig. 3**, **b)**, whereas the opposite trend can be seen for stiff hydrogels (25 kPa) (**Fig. 3**, **c**). Notably, Piezo1 knock down abrogates increased nuclear spreading area on soft V+ substrates as well as on stiff V- hydrogels (**Fig. 3**, **d**), suggesting that viscoelasticity and Piezo1-mediated cytoskeletal tension directly act on the nucleus^33^.

**Fig. 3.**
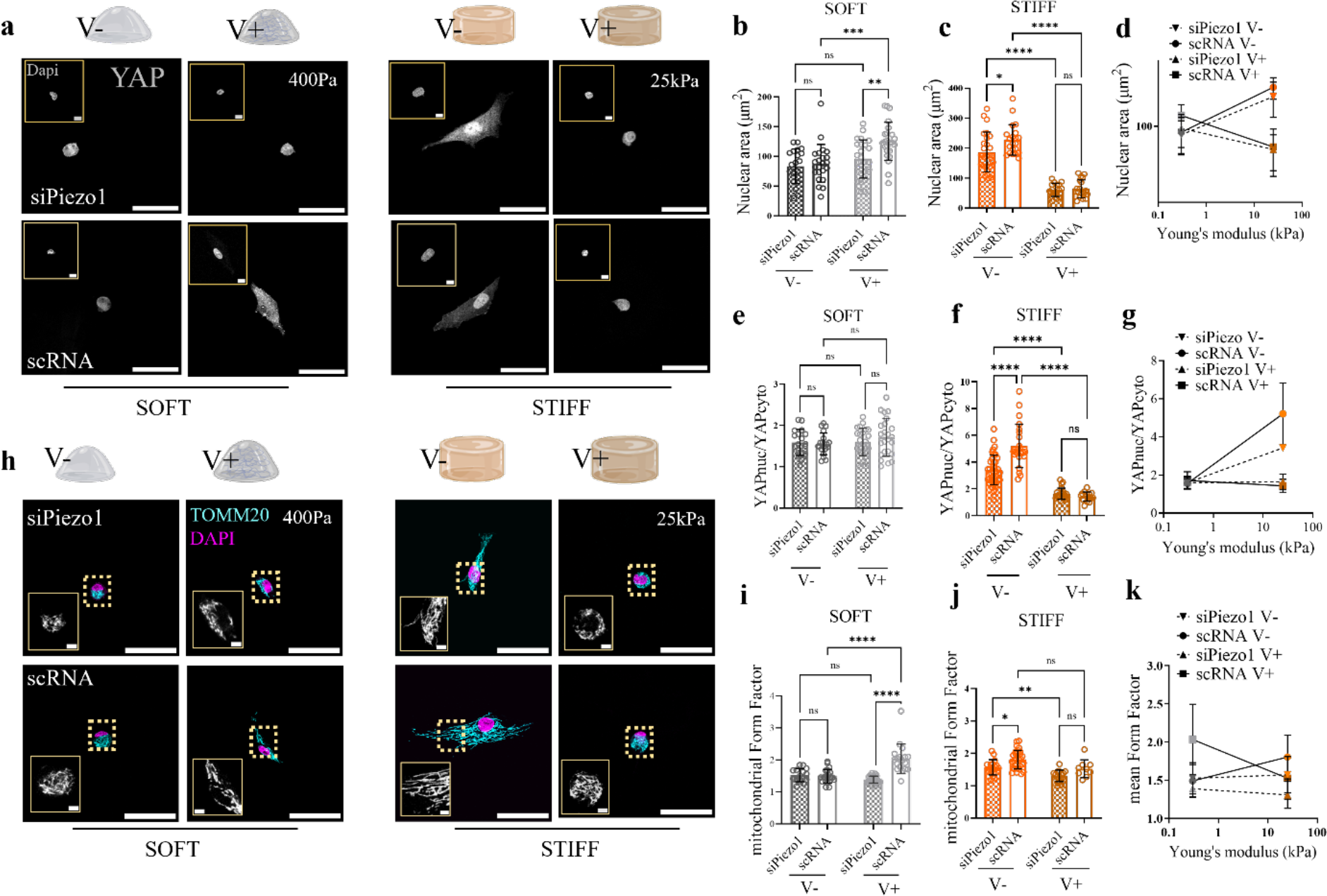
Matrix viscoelasticity and Piezo1 expression regulate downstream mechanotransduction and mitochondrial morphology. (**a**) Representative images of nuclei (insets) and YAP in siPiezo1 (top) and scRNA Y201 (bottom) cells on soft (left) and stiff (right) hydrogel groups of varying stress relaxation. Scale bar=50 µm, Dapi inset scale bar=5 µm. (**b, c**) Quantified nuclear spreading area on the soft (left) and stiff (right) hydrogel groups. All individual points represent individual nuclei measurements. Data shown as mean ± SD (n ≥ 16). (**d**) Summary of mean nuclear spreading area ± SD plotted as a function of Young’s modulus for all conditions. (**e, f**) Quantified nuclear over cytoplasmic YAP (nucYAP/cytoYAP) ratio on the soft (left) and stiff (right) hydrogel groups. All individual points represent individual cell measurements. Data shown as mean ± SD (n ≥ 15). (**g**) Summary of mean nucYAP/cytoYAP ratio ± SD plotted as a function of Young’s modulus for all conditions. (**h**) Representative images of siPiezo1 (top) and scRNA (bottom) Y201 MSCs immunostained for TOMM20 (cyan) and Dapi (magenta) cultured on soft (left) and stiff (right) hydrogels for 48h. Scale bar 50 µm; inset scale bar 2 µm (**i, j**) Quantified mean mitochondrial form factor of cells on the soft (left) and stiff (right) hydrogel groups. (All individual points represent individual cells mean mitochondrial form factor). Data shown as mean ± SD (n ≥ 9). (**k**) Summary of mean mitochondrial form factor ± SD plotted as a function of Young’s modulus for all conditions. (**b, c, e, f, i, j**) Statistical analyses were performed using a two-way ANOVA test. P values indicating significance, ns > 0.05, * ≤ 0.05, ** ≤ 0.01, *** ≤ 0.001, **** ≤ 0.0001.

In recent work by Pere Roca-Cusachs^34^, the idea that the force applied to the nucleus could dictate the nuclear translocation of important transcription factors such as Yes-associated protein (YAP), independently of other specific signalling pathways, was proposed. Authors described that nuclear flattening (i.e., increased projected spreading area) increased nuclear pore permeability via two mechanisms: partially opening nuclear pores to facilitate entry and by increasing nuclear membrane curvature. Previous work has reported the importance of Piezo1 in sensitively sensing tensional changes to regulate nuclear size in response to exogenously applied shear stress^35^. Thus, we sought to understand whether the observed morphological changes in the nucleus reflected transcription factor translocation by assessing the localisation of the mechanical rheostat YAP (**Fig. 3**, **a**). YAP translocated into the nucleus (YAPnuc/YAPcyto > 2) in response to increased substrate stiffness (**Fig. 3**, **e-g**), however, on soft substrates, YAP was mostly cytoplasmic and did not translocate in response to enhanced stress relaxation. YAP nuclear translocation in response to molecular clutch activation mechanisms has been shown to occur past an elasticity threshold of *E* ∼ 5 kPa^6^. Therefore, soft viscoelastic (soft V+) substrates do not activate molecular clutch mechanisms past this threshold. This prevents the nuclear translocation of YAP, despite cells demonstrating enhanced cell spreading, adhesion length, traction forces, decreased actin retrograde flow speed and increased nuclear projected area in soft viscoelastic (V+) compared to soft elastic (V-) substrates. Therefore, faster stress relaxation promotes mechanoactivation and mechanotransduction on soft substrates, although to a lesser extent when compared to stiff elastic matrices (stiff V-) eliciting the same effects.

Still, on stiff substrates, faster substrate relaxation (stiff V+) reduced both nuclear spreading area and YAP translocation. Additionally, Piezo1 activity has been linked to enhanced YAP nuclear translocation^17^. Indeed, on FN coated glass we observed that Piezo1 knock down slightly reduced the transcription factor nuclear translocation (**Supplementary Data Fig. 4**). The same relationship was observed on stiff elastic (V-) substrates, where siPiezo1 MSCs show significantly lower levels of nuclear YAP (**Fig. 3, c**).

We then hypothesised that cell-substrate interaction affected the cell’s metabolic capacity, as recent evidence linked stiffness-dependent integrin signalling to modulation of mitochondrial activity^36^. Oxygen consumption rate (OCR) experiments were preliminarily performed on *stiff* and *soft* Matrigel substrates (**Supplementary Data Fig. 5, a**). Matrigel substrates were used in this case due to the experimental implications of the Seahorse assay, which made the implementation of viscoelastic PAAm substrates into the system complex, whereas *soft* (*E*∼ 200Pa) and *stiff* (*E*∼ 1GPa) Matrigel systems have previously been implemented into the experimental set-up^37^. OCR measurements indicated that Y201 MSC mitochondrial respiration rates were sensitive to stiffness and Piezo1 expression (**Supplementary Data Fig. 5, b, c and d**), whereas non-mitochondrial respiration was only significantly decreased by knocking down Piezo1 on stiff Matrigel substrates (**Supplementary Data Fig. 5, e**). We found that a soft matrix would decrease cellular respiration capacity independently of Piezo1 expression (**Supplementary Data Fig. 5, a**). However, on stiff substrates, mitochondrial-dependent respiration would be increased, but this increase would be abrogated if Piezo1 was knocked down (**Supplementary Data Fig. 5, e**). Data thus proposed the mitochondria as a main sensor of matrix mechanics. Therefore, in order to assess mitochondrial respiration in response to varying viscoelasticity, we tagged the outer mitochondrial membrane protein TOMM20 (**Fig. 3**, **h**) and assessed mitochondrial elongation by quantifying the mean mitochondrial Form Factor in all experimental conditions (**Fig. 3**, **i-k**). Mitochondria appeared shorter on soft elastic (Soft V-) substrates compared to stiff elastic (Stiff V-) ones, as previously reported^36^. Increased stress relaxation on soft hydrogels promoted mitochondrial elongation and avoided matrix induced mitochondrial fission (**Fig. 3, i**). However, on stiff substrates, there were little changes in mitochondrial elongation induced by either increasing substrate relaxation or Piezo1 expression (**Fig. 3**, **j**). This is highlighted in **Fig. 3**, **h**, where mitochondrial elongation is mostly reduced in soft elastic (Soft V-) substrates, or when Piezo1 is knocked down across stiffnesses. Mitochondrial elongation provides information on the fusion vs. fission events of the mitochondrial network^38^, therefore highlighting the level of oxidative stress present in the cell. However, it is also worth quantifying the total mitochondrial area in the cell, to estimate respiration rates^39^. Indeed, from total mitochondrial area data **(Supplementary Data Fig. 6**), it is possible to hypothesise that overall cell respiration would be highly linked to mitochondrial mass inside the cell. This means that increased substrate relaxation (V+) recovers cellular respiration and metabolism in soft ECMs. Whereas on stiff matrices, increased stress relaxation decreases overall cellular respiration and slows the cell’s metabolic capabilities but does not induce a state of oxidative stress. Regarding the effect of Piezo1 in mitochondrial dynamics, it appears that channel knock down generally promotes mitochondrial fission (**Fig. 3, k**). This is most likely a consequence of the channel’s role in mediating the cell’s interaction with its environment. Thus, mitochondrial fission induced by Piezo1 knock down is a read-out of the effects of Piezo1 on overall cell morphology.

### Matrix viscoelasticity and Piezo1 influence transcriptomic phenotype

Finally, to monitor matrix viscoelasticity and Piezo1 activity-dependent transcriptional changes and obtain reference transcriptomic phenotypes, we performed RNAseq on all experimental conditions at the timepoint of previously shown mechanotransduction and metabolic data (48 h). This timepoint was chosen for two reasons, i) to compare RNAseq data with previously shown data; and ii) to agree with the timepoint of previous *omics* studies involving MSCs response to varying stiffness, stress relaxation and ligand density in 3D matrices, where authors performed RNAseq at 40 h to ensure formation of mature adhesions while minimising cell proliferation^40,41^.

Differential expression analysis was performed and a heatmap with up and down regulated genes was plotted for each hydrogel stiffness range **Fig. 4, a, and Fig. 5**, **a**). The total differentially expressed (DE) genes with a p value lower than 0.05 were used to obtain a visual representation of inner-group transcriptome variation. In the stiff group, a total of 731 genes were differentially expressed across conditions. Additionally, four distinct expression patterns emerge, which demarcate the four different experimental conditions assessed, highlighting both the roles of viscoelasticity and Piezo1 expression in regulating the cell’s transcriptome. Thus, to further interpret transcriptomic data, we performed gene ontology (GO) enrichment analysis on the DE genes from each experimental comparison. GO enrichment places top differentially expressed genes in functional modules of relevant sub-ontologies^42–44^. Firstly, we assessed the enrichment of the DE genes from scRNA cells seeded on stiff elastic (V-) versus viscoelastic (V+) matrices to describe the processes which drive Y201 MSCs’ response to viscoelasticity at this stiffness (**Fig. 4**, **b**). Top differentially expressed processes involved *cell motility, bone remodelling* and *G-protein-coupled receptor signaling* sub ontologies. These processes are in accordance with literature describing cell interaction with stiff 2D viscoelastic substrates. Indeed, our group has demonstrated that cell motility is reduced in epithelial cells in response to enhanced substrate stress relaxation at *E >* 1kPa^45^. Similarly, at stiffness *E* > 1kPa, we have shown that viscoelasticity reduces cell spreading area, molecular clutch engagement and YAP nuclear translocation, which are all processes involved in *bone remodelling.* In previous work by our group, it was demonstrated that substrate stress relaxation in hydrogels of similar stiffness (*E* ∼ 13 kPa) promoted chondrogenesis in hMSCs^46^. Finally, the *G protein-coupled receptor (GPCR) pathways* ontology is also highlighted. This involves genes belonging to the Ras-subfamily of GTPases (RASD1) or Rho signaling (GNAT1), which are processes that have been previously highlighted in studies that describe cell response to 2D viscoelastic substrates^47^.

**Fig. 4.**
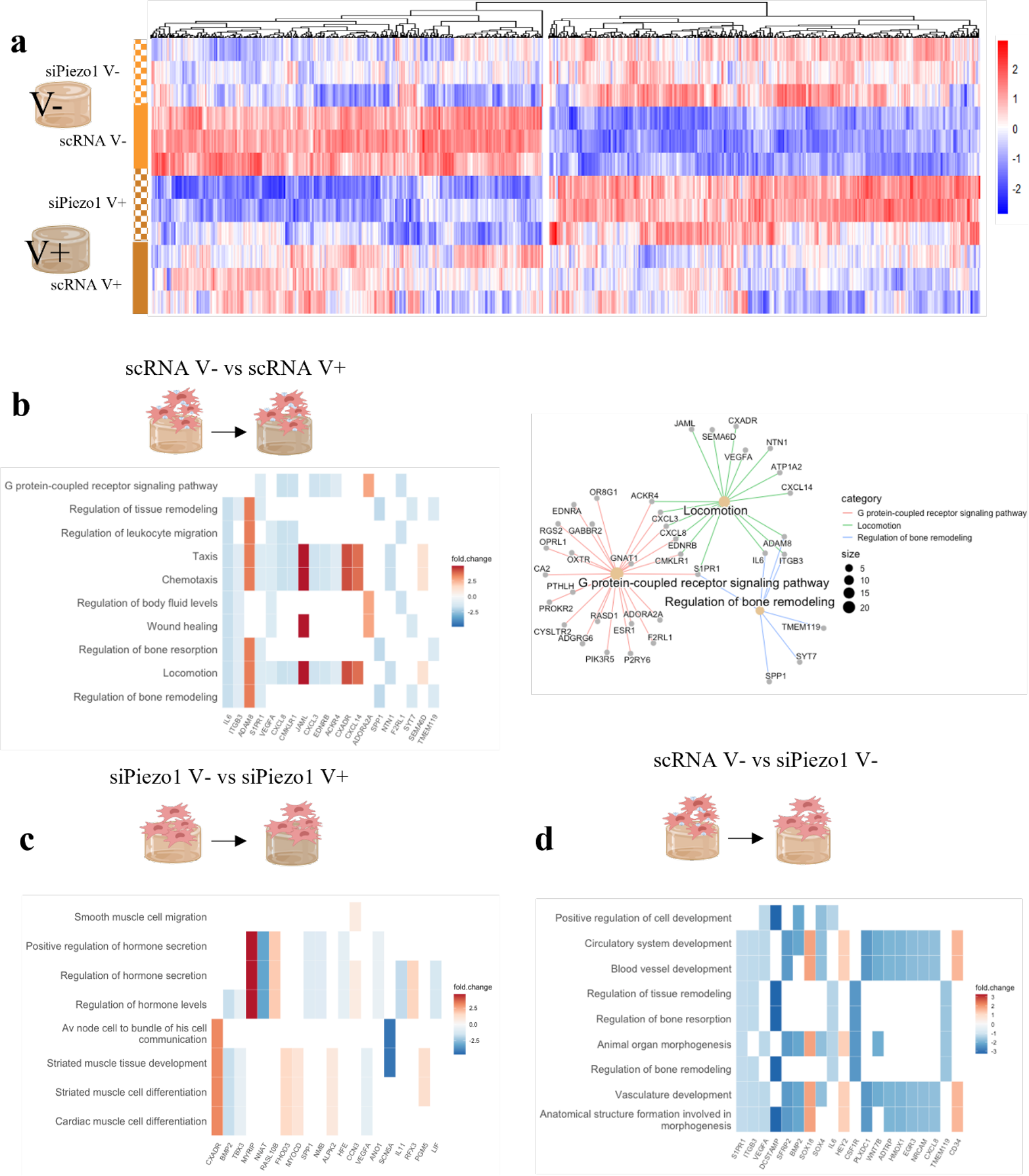
RNA-seq analysis of siPiezo1 and scRNA Y201 MSCs viscoelasticity sensing on stiff matrices. (**a**) Heatmap for Stiff group genes. A total of 731 genes are shown p < 0.05. (**b**) Heatmap of enriched results from Over Representation Analysis (ORA) overlapping most differentially expressed genes in the scRNA V- vs scRNA V+ comparison, this was accompanied by the enriched network of top enriched groups connected by overlapping genes (**c**) Heatmap of enriched results from ORA overlapping most differentially expressed genes in the siPiezo1 V- vs siPiezo1 V+ comparison (**d**) Heatmap of enriched results from ORA overlapping most differentially expressed genes in the scRNA V- vs siPiezo1 V- comparison.

**Fig. 5.**
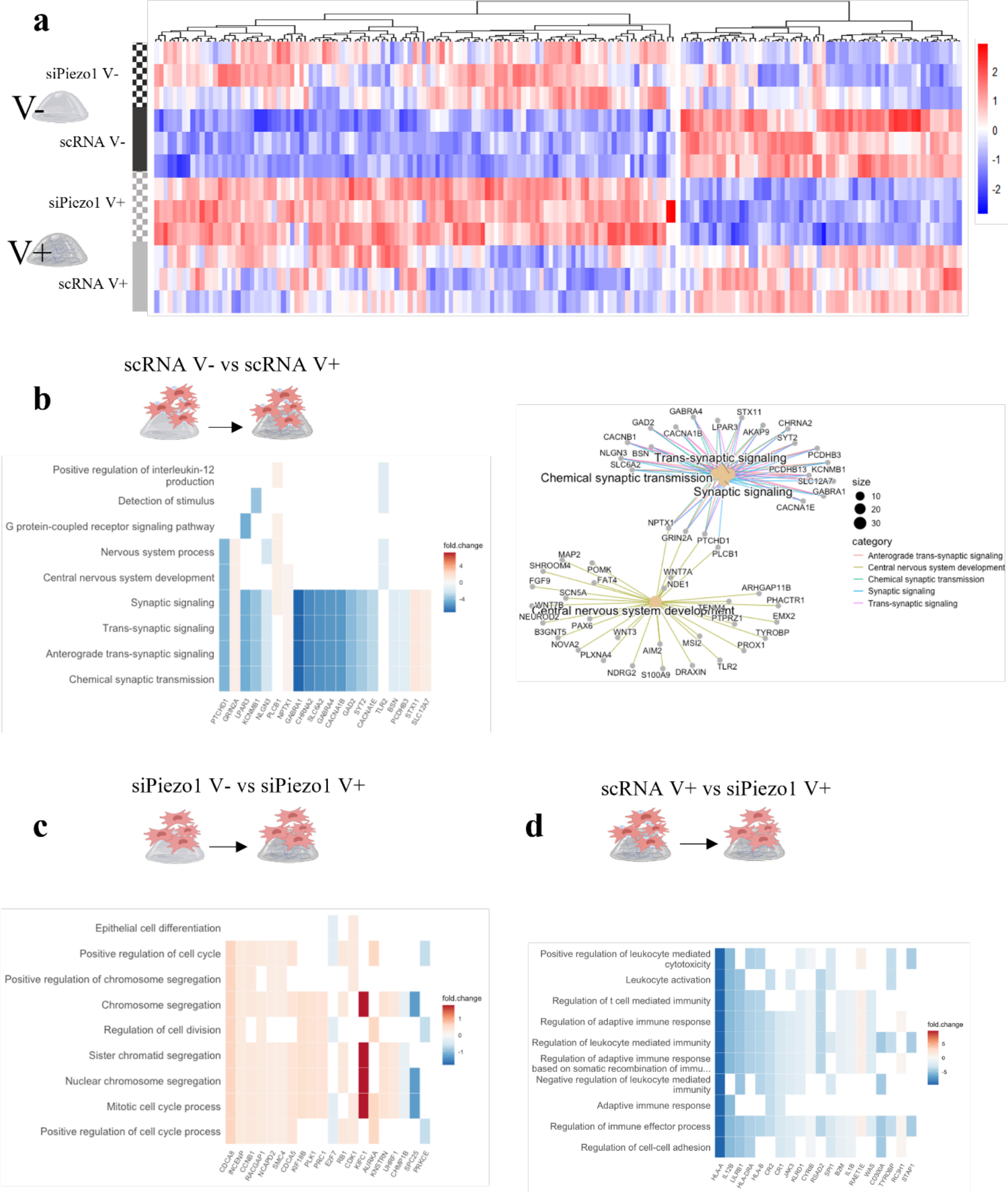
RNA-seq analysis of siPiezo1 and scRNA Y201 MSCs viscoelasticity sensing on soft matrices. (a) Heatmap for Soft group genes. A total of 177 genes are shown p < 0.05. **(b)** Heatmap of enriched results from Over Representation Analysis (ORA) overlapping most differentially expressed genes in the scRNA V- vs scRNA V+ comparison, this was accompanied by the enriched network of top enriched groups connected by overlapping genes (**c)** Heatmap of enriched results from ORA overlapping most differentially expressed genes in the siPiezo1 V- vs siPiezo1 V+ comparison **(d)** Heatmap of enriched results from ORA overlapping most differentially expressed genes in the scRNA V- vs siPiezo1 V- comparison.

This was followed by performing the same comparison on siPiezo1 cells to investigate Piezo1- independent transcription mechanisms of matrix viscoelasticity for this stiffness. In this comparison, processes such as *hormone secretion and regulation, muscle tissue development and differentiation* and *smooth muscle cell migration* were highlighted (**Fig. 4**, **c**). Data from this comparison allows us to hypothesise that Piezo1 expression is crucial for mediating viscoelasticity-induced changes in *bone remodelling* and *locomotion* processes. However, when Piezo1 is silenced in Y201 MSCs, enhanced substrate stress relaxation affects myosin related ontologies (i.e., *muscle tissue development and differentiation*). Our results thus suggest that myosin signalling mechanisms in the cell respond to enhanced viscoelasticity independently of Piezo1 expression in Y201 MSCs.

Finally, the comparison of scRNA vs siPiezo1 Y201 MSCs cultured on stiff elastic matrices was investigated. Here, top DE genes belonged to *cardiovascular system remodelling, bone remodelling* and *morphogenesis* processes. In this case, Piezo1 KD downregulates genes such as BMP2, VEGFA, ITGB3, which are involved in proliferation, differentiation, and integrin-related signalling in the cell. In fact, these processes have been directly associated with Piezo1 activity and expression in MSCs^48^, and agree with data shown in this work, where we have demonstrated that Piezo1 knock down reduces overall molecular clutch engagement (**Fig. 2**, **c, g** **and k**) on stiff elastic (stiff V-) matrices and glass substrates (**Supplementary Data Fig. 2**) as well as YAP nuclear translocation (**Fig. 3**, **f**, **Supplementary Data Fig. 4)**, which has been shown to promote bone differentiation in hMSCs^49^.

The same analysis was performed for cells cultured on the soft group. In this case, 177 genes were differentially expressed (**Fig. 5, a**), which suggests that at this stiffness range, both Piezo1 expression and substrate stress relaxation infer a lesser effect on the cell’s transcriptome. Still, four distinct expression patterns emerge in the plotted heatmap (p < 0.05), which correspond to the four different experimental conditions. We then assessed enriched DE genes in scRNA cultured on soft elastic (soft V-) versus viscoelastic (soft V+) matrices **Fig. 5, b**). In this comparison, processes such as *GPCR pathway, synaptic signaling* and *Central nervous system development* were featured. Interestingly, in the *Central nervous system development* subontology we find genes that belong to the Wnt family (WNT7A, WNT3), which have been described as important regulators of YAP and have been found to respond to matrix stiffness^50^. Whereas we found that at this stiffness range (∼400Pa), viscoelasticity did not promote YAP nuclear translocation in cells (**Fig. 3**, **e**), it is possible that precursors such as WNT3 and GPCRs (Gα subunits) belonging to non-canonical Wnt signaling^51^ still become activated to support cell proliferation in response to enhanced substrate stress relaxation. These ontologies have also been highlighted in previous studies that assessed transcriptional changes in MSCs encapsulated in dynamic viscoplastic matrices when compared to fully crosslinked *elastic* ones^52^. Additionally, we found that *synaptic signaling* processes were underscored in the enrichment analysis. Indeed, genes in this subontology include phospholipase C beta 1 (PLCB1), which has been associated to cytoskeletal rearrangement processes in gastric tumour tissue samples^53^ as well as other genes which encode for Ca^2+^ responsive channels (CACNA1E and 1B, KCNMB1). These results could potentially explain our data regarding enhanced cell spreading area (cytoskeletal rearrangement) and clutch engagement (**Fig. 2**, **f, and** **Fig. 2, b, f, j**) in scRNA cells in response to increased viscoelasticity on soft matrices.

This was again followed by a comparison of siPiezo1 cells in soft elastic (V-) vs soft viscoelastic (V+) matrices to highlight the Piezo1-dependent cell response to viscoelasticity as this stiffness (**Fig. 5**, **c**). Strikingly, enrichment analysis demonstrated that most of the previously shown GOs were replaced by *cell division* processes, which were mostly up regulated in viscoelastic hydrogels in a Piezo1 independent way. In our data, we do not see any phenotypic difference between siPiezo1 MSCs cultured on soft elastic (V-) vs viscoelastic (V+) hydrogels. Nonetheless, it may be that viscoelasticity in soft matrices promotes cell proliferation, as previously reported in a cancer cell line^9^, and this may happen independently of Piezo1.

Finally, we compared scRNA and siPiezo1 cells on soft viscoelastic (V+) hydrogels (**Fig. 5**, **d**). Surprisingly, Piezo1 knock down promoted the down regulation of several ontologies relating to *immune response regulation* and *cell-cell adhesion*. MSCs have important immunomodulatory properties, which have been shown to be responsive to matrix mechanics^54,55^. Similarly, Piezo1 mechanosensing has recently been linked to adherens cell-cell junction formation in endothelial cells^56,57^ as well as immune cell migration^58^, however, the link between Piezo1 knock down and MSC downregulation of immunomodulatory genes remains unclear, and something that should be addressed in future work. Interestingly, in previous work assessing transcriptomic changes in MSCs encapsulated in 3D hydrogels of increasing stiffness (*E* increasing from 3 kPa to 18 kPa), several immunomodulatory markers were differentially expressed^40^. This was found to be linked to stiffness induced activation of the immune and inflammatory transcription factor NFkB-p65. Our data therefore proposes Piezo1 as a mediator of stiffness-induced immunomodulation in MSCs, which occurs at softer regimes.

Overall, transcription studies evidenced how both viscoelasticity and Piezo1 activity regulate a diverse range of processes in MSCs. To expand on molecular clutch and mechanotransduction data previously shown in this study, we have also demonstrated that depending on the stiffness range, different gene groups respond to changes in matrix viscoelasticity as well as Piezo1 knock down.

## Conclusions

This study introduces a platform in which to study MSC substrate interaction in response to both viscoelasticity and elasticity. Previous studies have shown that cells respond to matrix viscoelasticity via dynamic molecular clutch mechanisms that sense and engage with the cell’s mechanical microenvironment in a stiffness-dependent way^9,28^. Here, we demonstrate that Piezo1 is an important sensor of matrix viscoelasticity as knocking down the channel in the MSC cell line Y201 results in a mechanobiologically impaired phenotype with reduced cell spreading behaviour, molecular clutch engagement, mechanotransduction and mitochondrial metabolic activity. In addition, we characterise how cells respond to viscoelasticity in a stiffness-dependent manner, where on soft (*E* ∼ 400 Pa) matrices, faster stress relaxation promotes cell mechanoactivation and on stiff (*E ∼* 25 kPa) substrates, the opposite is true^9^. Most importantly, we demonstrated that cell mechanoactivation in response to faster stress relaxation in soft matrices is abrogated when Piezo1 is knocked down, highlighting the role of the channel in relaying viscoelasticity at this stiffness. Finally, by performing RNAseq, we obtain transcriptomic phenotypes of cell-substrate interaction, obtaining gene signatures that describe how cells respond to substrate viscoelasticity at two stiffnesses and in terms of Piezo1 expression.

Our findings expand on our current understanding of cell response to substrate viscoelasticity and place the mechanosensor Piezo1 as a key mediator of these cues in soft regimes. Soft tissues in the body showcase the highest degree of viscoelasticity such as brain, lung tissue or bone marrow^8^. Coincidentally, Piezo1 has been implicated in the physiological regulation and pathophysiology of the central nervous system, being originally identified in a mouse neuroblastoma cell line^12^, as well as being shown to regulate the differentiation of neural pluripotent stem cells^17^. Indeed, many of the differentially expressed genes in the soft group comparison (which emulates the stiffness of native brain^59^) highlighted genes inherently linked with the central nervous system and synaptic transmission processes (**Fig. 5**, **b**).

While our study was conducted in an MSC line, it opens the scope to investigate how viscoelasticity and Piezo1 mediate cell behaviour in a more specific physiological context such as neural tissue. Here, soft viscoelastic matrices might provide a suitable platform in to further our understanding of tissue physiology, and different tools ranging from molecular cell-ECM interaction studies, mechanotransduction and transcriptomics can be employed to address tissue-specific questions.

## Materials and Methods

### PAAm hydrogel synthesis

Hydrogels were polymerised on clean borosilicate 12 mm diameter glass coverslips (VWR). Coverslip surfaces were functionalised with 3-(Acryloyloxy)propyltrimethoxysilane (Alfa Aesar). Hydrogel solutions prepared using stock solutions of 40% Aam (sigma) and 2% bisAam (sigma) mixed in different ratios for each hydrogel composition (**Supplementary Data Table 1**). For hydrogels requiring the addition of linear acrylamide, this was prepared beforehand by mixing 50ul of 40% acrylamide in 317 µl dH20 and polymerising it with 25 µl 1.5% TEMED and 8 µl 5% APS for 2h at 37°, making a final concentration of 5% linear acrylamide. Once mixed, all hydrogel solutions were thoroughly mixed and vortexed prior to use. A hydrogel solution drop of 12 µl was placed on top of a hydrophobic (RainX treated) coverslip and an (acryloyloxy)propyltrimethoxysilane treated coverslip was placed on top of the drop to synthesise flat hydrogels. Gelation was allowed to occur at RT for 30 min before detaching and swelling in PBS overnight at 4°C.

To promote cell adhesion, substrates were functionalised with full length fibronectin. This was done by placing 0.2 mg/ml sulfo-succinimidyl-6-(4-azido-2-nitrophenyl-amino) hexanoate (sulfo-SANPAH) (Thermo Fisher) in 0.5 mM pH 8.5 HEPES buffer onto the hydrogel surface and irradiating with ultraviolet (UV) light (365 nm) at a distance of 3 inches for 20 min. The darkened sulfo-SANPAH solution was removed, and substrates were rinsed twice with HEPES buffer and incubated with 10 µg/ml of fibronectin in HEPES at 37°C, overnight. All substrates were exposed to UV light in a sterile culture hood for 30 min. prior to use. Before plating cells, hydrogels were equilibrated in cell culture medium for 30 min. at 37°C.

### Mechanical characterisation of hydrogels

Nanoindentation measurements were performed using a nanoindentation device (Chiaro, Optics11 Life) adapting a previously reported approach ^60^.

Measurements were performed at RT in PBS unless stated otherwise. To obtain the Young’s modulus of the substrates, single indentation curves (n > 75) were acquired at a speed of 2 µm/s over a vertical range of 10 µm, changing the (x, y) point at every indentation. The selected cantilever had a stiffness of 0.52 N/m and held a spherical tip of 27.5 µm radius. A minimum of three maps per replicate were measured. All collected curves were pre-processed and analysed with a custom-made graphical user interface, the analysis and all software used are described in detail in ^60^.

To perform stress relaxation measurements, ≥ 95 indentations were performed, each spaced at least 50 μm from the previous. The selected cantilever had a stiffness of 0.52 N/m and held a spherical tip of 27.5 µm radius. For each indentation, the probe moved at a strain rate of 5 μms^-1^ until it reached an indentation depth (h) of 3 μm, which was maintained for 60 s using the instrument’s closed feedback *Indentation control mode*. The applied strain (!_!_) for the stress relaxation measurements was calculated as: ! = 0.2 ∗ (/*, where (= √ℎ ∗ *, where h is the indentation depth and R the probe radius ^61^.

Therefore, with an indentation depth of 3 μm and a probe radius of 27.5 µm, the applied strain was approximately 7%.

Acquired data was pre-processed using a previously published open-source software (time branch of the project) ^62^. To analyse the stress relaxation behaviour of the material, an analysis script in the form of a jupyter notebook was developed ^63^. Briefly, force-time F(t) curves were first aligned to zero force if their baseline was negative. Then, the maximum of F(t) and its corresponding time was found, yielding the point (t0, F0). Curves were therefore aligned to 0 time by a horizontal shift equal to t0. Following this, the signal was cropped between t0 and the maximum time before retraction, i.e., only the part of the signal where the indentation was kept constant was retained. Following this, F(t) was normalised by dividing the whole signal by F0. Because individual curves were too noisy to be analysed, an average curve was found and used for quantification of the time for the stress to decrease to an 80% of the original value as well as the energy dissipation of the materials. This was done by extracting the time at which force reaches 80% of original value.

To perform rheology measurements, hydrogels were prepared in 15 mm diameter PDMS moulds using 250 µl volumes. Samples were left to swell overnight and subsequently measured with a Physica MCR 301 rheometer (Anton Parr). The linear viscoelastic region was determined by performing amplitude sweeps from 0.01 to 10 % stain and then a strain of 0.1% was used to obtain frequency sweeps from 100 to 1 rad/s.

### Y201 hMSCs culture and transfection

Y201 hMSCs were kindly donated by Professor Paul Genever from the University of York. Cells were grown in as adherent cultures in high glucose DMEM (Gibco), 10% FBS and 1% P/S. Cells were passaged every three days and used between passages 70-90.

The Piezo1 channel was routinely knocked down prior to experiments using small interfering (si) RNA. This was done with the pre-validated siRNAs library from Thermo Fisher. Specifically, siRNA (5’ -> 3’ GCUUCACGUUUUCAAGCUGtt and 3’ -> 5’ CAGCUUGAAAACGUGAAGCtt). A validated control siRNA Ambion™ Silencer™ Negative Control #1 was also used to ensure transfection efficiency and validate the channel knock down (scRNA). All siRNA was resuspended in Nuclease Free water (provided with kit) at 100 µM and stored at -80°C. Prior to transfection, cells were plated on cell culture treated six well plates. Cells were left to adhere in antibiotic free supplemented media. After adhering for 24h, media was changed to OptiMEM (Gibco) reduced serum medium and siRNA was introduced with Lipofectamine RNAiMax transfection reagent (Thermo Fisher), as per manufacturer’s instructions. Briefly, 25pMol of siRNA/scRNA and 7.5 µl of RNAiMax lipofectamine were used per well and incubated for 72h. After incubation, media was changed to supplemented high glucose DMEM (Gibco) and cells were left to recover overnight prior to use for experiments.

For performing actin flow experiments, cells were transfected with a LifeAct-GFP plasmid (Ibidi) using a neon electroporator system (Invitrogen). 5 µg plasmid were used per 100 µl electroporator tip, as per manufacturer’s instruction.

### RNA extraction

RNA extraction was performed using a commercially available kit (RNease Micro kit, Qiagen). RNA concentration and quality was monitored by spectrophotometry using the Nanodrop 2000 (Thermo Scientific).

### RT-qPCR

For quantifying gene expression, real time quantitative Polymerase Chain Reaction was performed (RT-qPCR). 300-500ng of RNA was used to generate complementary DNA (cDNA) using QuantiTect Reverse Transcription Kit Reagents (Qiagen). cDNA was amplified using Quantifast SYBR green qPCR kit (Qiagen) with specific primers for Piezo1 and ribosomal protein L3 (RPL3), which was used as a genetic internal control. Expression was quantified using the 2-ΔΔCt method and amplification was carried out using an Applied Biosystems 7500 Real Time PCR system (Thermo Fisher).

### In cell western

For In-cell western Piezo1 protein quantification cells were seeded on a 48 well plate and cultured for 24h post siRNA transfection. Three cell conditions were assessed: scRNA control, siPiezo1 and untransfected Y201 MSCs. After 24h, cells were fixed for 15 min with 4% paraformaldehyde with 1% sucrose and subsequently permeabilised with permeabilising buffer (10.3 g Sucrose, 0.292 g NaCl, 0.06 g MgCl2 (hexahydrate), 0.476 g HEPES and 0.5 mL Triton-X100, in 100mL, pH 7.2). Samples were washed with 3x with PBS- and blocked with a 1% milk protein PBS solution for 1.5 h on an orbital shaker at room temperature. The primary antibody was incubated in blocking solution overnight at 4°C. The next day, samples were washed 3x with PBS and the secondary antibody was added at a dilution of 1:800 in blocking buffer (1:500 in the case of the CellTag for normalising the protein signal). Samples were then washed 5x with PBS and dried overnight in a chemical flow hood. Protein signal was measured with an Odissey Scanner at 700 and 800 nm once samples were dried. The intensity of the wells was then normalised to the cell-tag intensity and then normalised to the intensity of the samples which were only stained with the secondary antibody, to correct for background signal.

### Immunofluorescence staining

Samples were fixed with 4% formaldehyde in PBS for 15 min at room temperature (RT). 0.1% Triton X-100 in PBS was used to permeabilise cells for 10 min at RT. Samples were blocked in 1% BSA in PBS and incubated for 1h and then the primary antibody was either incubated for 1h at RT or overnight at 4°C in an incubation chamber. Samples were washed with 0.1% Triton X-100 in PBS and the secondary antibody was incubated in 1% BSA in PBS for 1h at RT. Finally, samples were washed thrice with 0.1% Triton X-100 in PBS and once in PBS. For image acquisition, samples were mounted on glass slides with Vectashield Hardset Antifade mounting medium with dapi.

See **Supplementary Data Table 2** for antibody dilution and manufacturer.

### Actin Flow measurements

On the day of imaging, LifeAct-GFP transfected cells were cultured on of the desired surfaces and left to adhere for a minimum of two hours. Cells were then placed on the heated stage of the LSM980 Zeiss confocal fluorescent system. Imaging was performed at 37°C in CO2 independent media (Gibco) with a 40x oil immersion objective with a numerical aperture of 1.3. Cells were illuminated with a 488 laser and images were acquired at a frame rate of 1 image every two seconds for a total of 4 min.

### Traction Force microscopy

Cells traction forces were measured using the EVOS M700 (Thermo Fisher) imaging system at 20X magnification. Cells were seeded on glass petri-bound hydrogels prepared with 1 µl/ml of 0.1% FluoSpheres Carboxylate-modified microspheres (0.2 µm, 580/605, 2%) (Thermo Fisher) aqueous solution. In each sample, a total of 6 positions were tracked and z-stacks were taken (an image was taken every 5µm) of both the cells and beads channels. Then, cells were detached with Trypsin for 10 min. and the same positions were imaged without cells to obtain the reference image.

### Oxygen Consumption Rate (OCR) measurements

OCR measurements were performed with the Agilent Seahorse XF Cell Mito Stress Test kit with an XF24 extracellular flux analyser (Seahorse Bioscience). XF24 plates were coated with either 23 µl of 100% Matrigel (soft) or 100 µl of 2% Matrigel (stiff). The solutions were evenly distributed with a flat pipette tip and the Matrigel was left to gel for 10 min. at 37 °C. 30,000 cells were seeded per well 24h before performing measurements. The experiment was conducted according to the manufacturer’s instructions with at least 4 technical replicates per experiment.

### RNA sequencing

RNA-Seq was performed by Glasgow Polyomics at the University of Glasgow. Briefly, strand-specific RNA-Seq libraries were prepared using the Illumina Stranded mRNA (poly A selected) library preparation kit with a NextSeq2000 system. In total, 100x100 bp paired end reads and an average of 30M total reads were generated for each sample. All libraries were aligned to the Homo- sapiens.GrCh.cdna genome using Kallisto 0.46.1^64^. To perform differential gene expression, aligned reads were imported to the DESeq ^65^ bioconductor package for R studio with the Txtimport function. Then, differential expression analysis was performed, and the counts matrix was obtained. Heatmaps of differential gene expression were obtained with the pheatmap function. GO terms analysis and graphs were obtained with the Interactive Enrichment Analysis tools implemented by the Gladstone Institutes Bioinformatics Core^44^.

### Image Analysis

#### Fibronectin functionalisation analysis

Immunofluorescent fibronectin hydrogel images were opened using Image J 2.14.0v (National Institute of Health, US). A square region of interest (ROI) was measured on three parts of the image, for all hydrogel conditions. A negative control (immunostained hydrogel without any fibronectin functionalisation) was used to normalise background signal. The reported fibronectin intensity values were therefore obtained by measuring the average integrated density per sample minus the negative control integrated density.

#### Cell/nuclear morphology analysis

Actin cytoskeleton images were converted to 8-bit, background was subtracted (rolling radius = 300) and a Gaussian blur of sigma 1 was applied. After this pre-processing, images were thresholded using Otsu’s method. Thresholded features were selected with the Wand function and the resulting ROIs were measured to quantify parameters such as cell and nuclear area and circularity.

#### Focal Adhesion analysis

Focal adhesion quantification was performed with an adapted version of the Horzum protocol, previously described in literature ^66^. Briefly, the vinculin channel images were cropped so that individual cells were in each image to analyse. Images were converted to 8-bit and background was subtracted (rolling radius = 50). Then, a Gaussian Blur (sigma = 1) was applied, and the contrast was enhanced with the CLAHE plugin (blocksize = 19, histogram = 256, maximum =3, mask = None), the mathematical exponential was applied to further minimise background and the brightness and contrast was adjusted automatically (saturated = 0.35). The log 3D filter was applied (sigmax = sigmay = 3), the LUT was inverted, and the image was thresholded using the Triangle method. The scale was adjusted according to the image pixel size calibration, adhesions were split with a watershed algorithm and finally the particles were analysed. Only particles over 0.75 µm^2^ were measured. The results were saved for individual adhesions as well as a summary of all particles in the cell.

#### Actin Flow analysis

Actin retrograde flow speed was calculated by kymograph analysis. In summary, timelapses were loaded onto image J and the area at the cell edge was sliced, producing a kymograph of line width 1 which plotted displacement over time using the Multi kymograph function. Flow speed was calculated by measuring and diving the bounding rectangle parameters (width over length) and converting it to nm units from pixel units.

#### Traction Force Microscopy analysis

Cell-generated traction forces were quantified using Image J 2.14.0v (National Institute of Health, US) and several plug-ins available open-source, following the protocol developed by Qingzon Tseng^67,68^. First, images were loaded onto ImageJ and stacked per channel, resulting in three different stacks: beads pre cell detachment (before), beads post cell detachment (after) and brightfield cell images. A maximum z-projection was created of the area closest to the cell-bead interface and saved. Then, the before and after images were opened and aligned with the Template Matching plugin. After the images were aligned, the displacement of the beads between the before and after pictures was calculated with the Particle Image Velocimetry (PIV) plugin. From PIV, a data file of displacement field vectors is obtained, which can be transformed into traction forces with the Fourier Transform Traction Cytometry (FTTC) plugin. With the FTTC plugin, the pixel size was scaled to microns using the appropriate conversion. Furthermore, the Poisson ratio and Young’s modulus of the hydrogels were specified per hydrogel in order to accurately calculate the forces exerted by the cells. Poisson ratio was always input as 0.5 and the Young’s modulus of each gel family was input as 400Pa (soft) and 25KPa (stiff). The average forces of each individual cell were summed (excluding those below 0.5 Pa) and plotted. Stress maps were created with the ParaView (v5.8.0) software, to visualise the force distribution in the cells.

#### YAP nuclear localisation analysis

The nuclear to cytoplasmic YAP ratio was determined as follows:

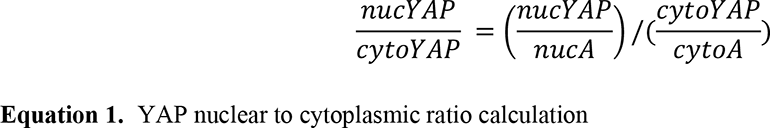

Where nucYAP is the integrated density of the YAP channel in the nucleus, nucA, the area of the nucleus (obtained with the Dapi channel); cytoYAP the integrated density of YAP in the cytoplasm, calculated as: cytoYAP = cellYAP – nucYAP (being cellYAP the integrated density of YAP in the cell as defined by the actin cytoskeleton channel. Similarly, cytoA is the area of the cell cytoplasm, defined as: cytoA = cellA – nucA, where cellA is the total cell area calculated from the actin cytoskeleton channel.

#### Mitochondrial morphology analysis

The Mitochondrial Analyzer plugin was opened and the 2D threshold options were optimised according to the quality of the images obtained. The Thresholding was optimised and performed as follows: background was subtracted (rolling (microns) = 1); a sigma filter plus was applied to further reduce background noise and smooth object signal (radius = 0.5, 2.0 sigma); contrast was enhanced with ‘enhance local contrast’ (max slope = 2) and gamma was adjusted (value = 0.8) to correct remaining dim areas. The threshold method employed was weighted mean with a block size of 1.25 µm and the C value was between 5 and 7 (this required adjustment from one image set to another, but it was empirically determined with the ‘2D threshold optimize’ menu available in the plugin). Finally, the 2D threshold menu has a few post-processing command options, of these, remove outliers (radius = 0.5 pixels) and show comparison of threshold to original were ticked. After a binary image of the mitochondria was obtained, the 2D analysis menu was clicked. Here, the per-cell analysis was performed as well as the per-mito analysis.

### Statistical analysis

Statistical analysis and graph plots were performed using GraphPad version 9.0.0 and R studio software. Unless stated otherwise, when two populations were contrasted, a t-test was performed. In the case of normal data distribution, unpaired t-test with Welch’s correction was performed; if data was not normally distributed, a Mann-Whitney t-test was performed. Normal distribution was assessed with D’Agostino Pearson normality tests. To compare several groups i.e., data per hydrogel pair, a two-way ANOVA was performed with Tukey multiple comparison correction. Data was shown as individual values in mean ± SD column graphs. P values indicating significance, ns > 0.05, * ≤ 0.05, ** ≤ 0.01, *** ≤ 0.001, **** ≤ 0.0001.

## Acknowledgements

We thank the Gladstone Bioinformatics Core for their interactive enrichment analysis tools. M.A.G.O acknowledges the Wellcome grant [204820/Z/16/Z], which supported the RNA sequencing shown in this work. M.S-S is grateful for financial support from the European Research Council AdG (Devise, 101054728) and EPSRC HT2050 grant (EP/X033554/1). IBEC is member of CERCA Programme / Generalitat de Catalunya. S.D. acknowledges a Worldwide Cancer Research grant 21-0156, AIRC Foundation investigator grants 21392 and 28940, and Italian Ministry of University and Research PRIN grants 2022T9RM8A and P2022CE7SP. P.R. acknowledges 2020, 2021, and 2022 Veronesi Foundation Postdoctoral Fellowships and an AIRC MFAG 27453.

## Author Contributions

M.A.G.O., M.V., M.S.-S. conceived the project. O.D., M.V. and M.S.-S. supervised the project. M.A.G.O., performed all experiments and analysed all data. P.R. and S.D. supervised the experiments performed in the University of Padova. G.C. and J.V. performed mechanical characterisation and data analysis together with M.A.G.O. J.L. developed and shared the protocol for the synthesis of Soft V- and Soft V+ matrices. M.A.G.O. wrote the first original draft of the article, which was reviewed and edited by G.C. and M.S.-S. The article was read and corrected by all authors, who contributed to the interpretation of results. Funding was acquired by M.S.-S., S.D. and M.A.G.O.

## Competing Interests

Authors declare no competing interests.

## Notes

### Competing Interest Statement

The authors have declared no competing interest.

