## Supplementary Data for "Piezo1 is a mechanosensor of soft matrix viscoelasticity"

### Supplementary Figures

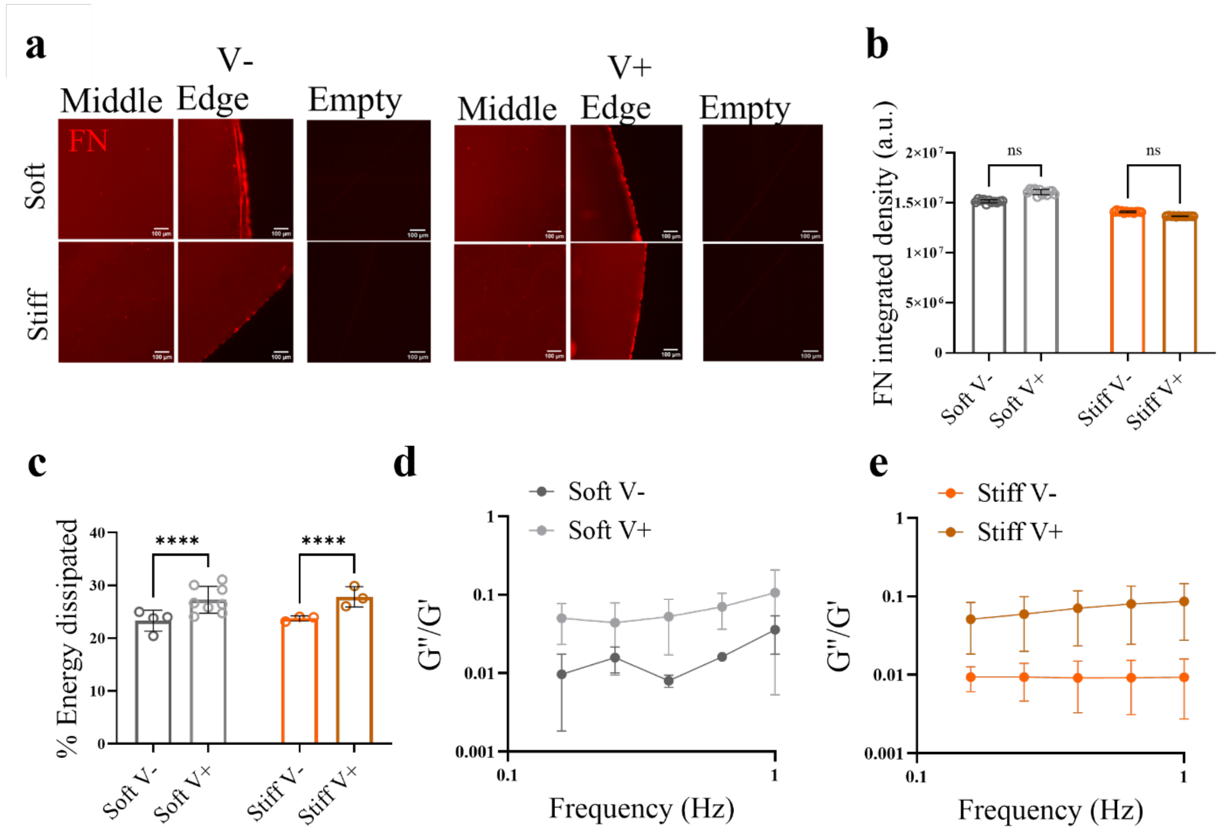


**Supplementary Fig. 1**. (**a**) Representative immunofluorescence images of the 10 µg/ml fibronectin coating (red) of the hydrogels, showing the middle and edge of the hydrogel on the glass coverslip. Images of stained non-functionalised hydrogels are also shown in the empty column, for each hydrogel type. Scale bar 100 µm. (**b**) Quantification of the resulting fibronectin staining intensity (integrated density a.u.) shown as mean ± SD of different areas of N>3 of each hydrogel. (**c**) Average energy dissipated in soft (grey) and stiff (orange) hydrogel groups, shown as mean ± SD of n ≥ 3 indentation maps on N ≥ 4 hydrogels. (**d**) G''/G' plot of bulk rheology frequency sweep measurements performed at 0.1% strain for the soft group (**e**) G''/G' plot of bulk rheology frequency sweep measurements performed at 0.1% strain for the stiff group. Shown as mean ± SD of N ≥ 3 hydrogels. P values indicating significance, ns > 0.05, **** ≤ 0.0001.

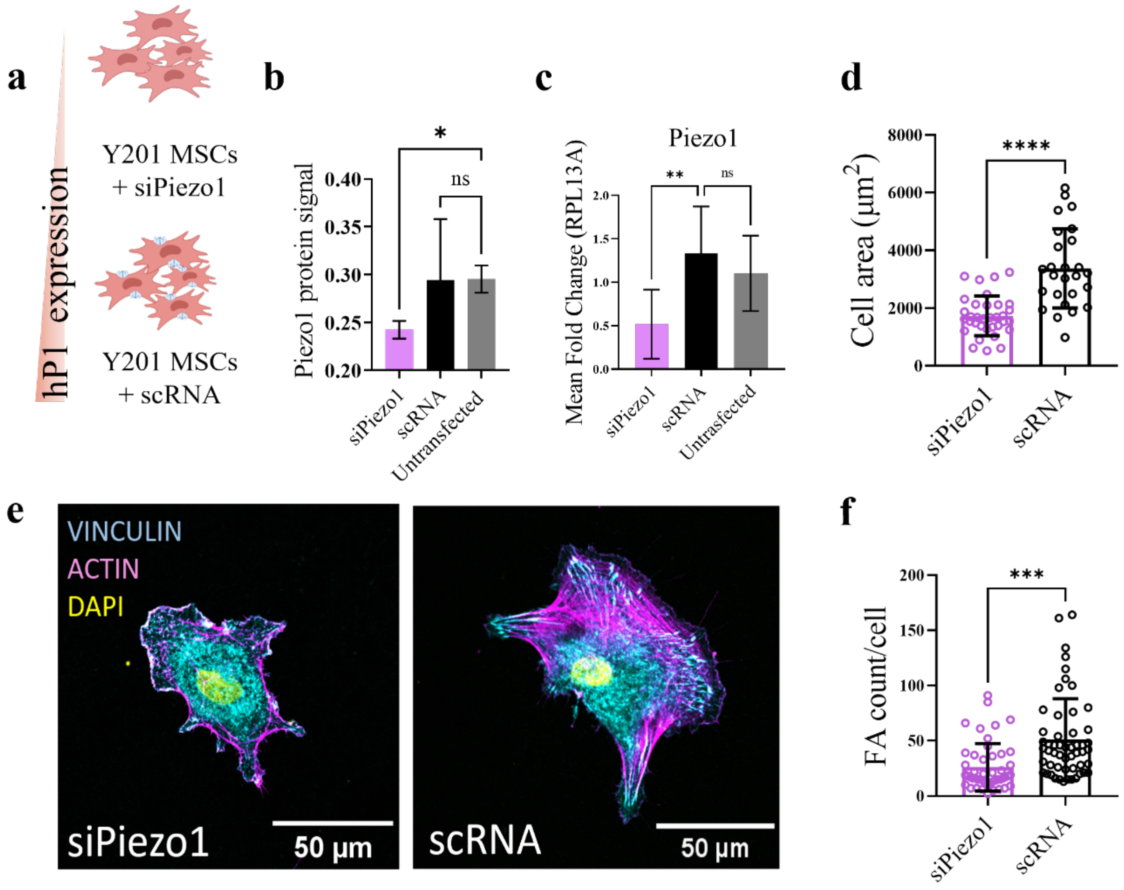


**Supplementary Fig. 2.** (**a**) Diagram describing the two treatment conditions: Piezo1 siRNA or scRNA was used on Y201 MSCs prior to performing experiments. (**b**) Piezo1 protein signal as quantified from an in cell western, shown as mean ± SD, n = 3. (**c**) Mean fold change of Piezo1 gene expression normalised to RPL13A, shown as mean ± SD, N ≥ 4. (**d**) Cellular area was quantified after cells were left to adhere for 48h, shown as individual cell values plus mean ± SD, n ≥ 26. (**e**) Representative images of cells cultured on fibronectin coated coverslips, scale bar 50 µm. (**f**) Number of focal adhesions per cell plotted as mean ± SD per individual cell, n ≥ 48. P values indicating significance, ns > 0.05, *** ≤ 0.001, **** ≤ 0.0001.

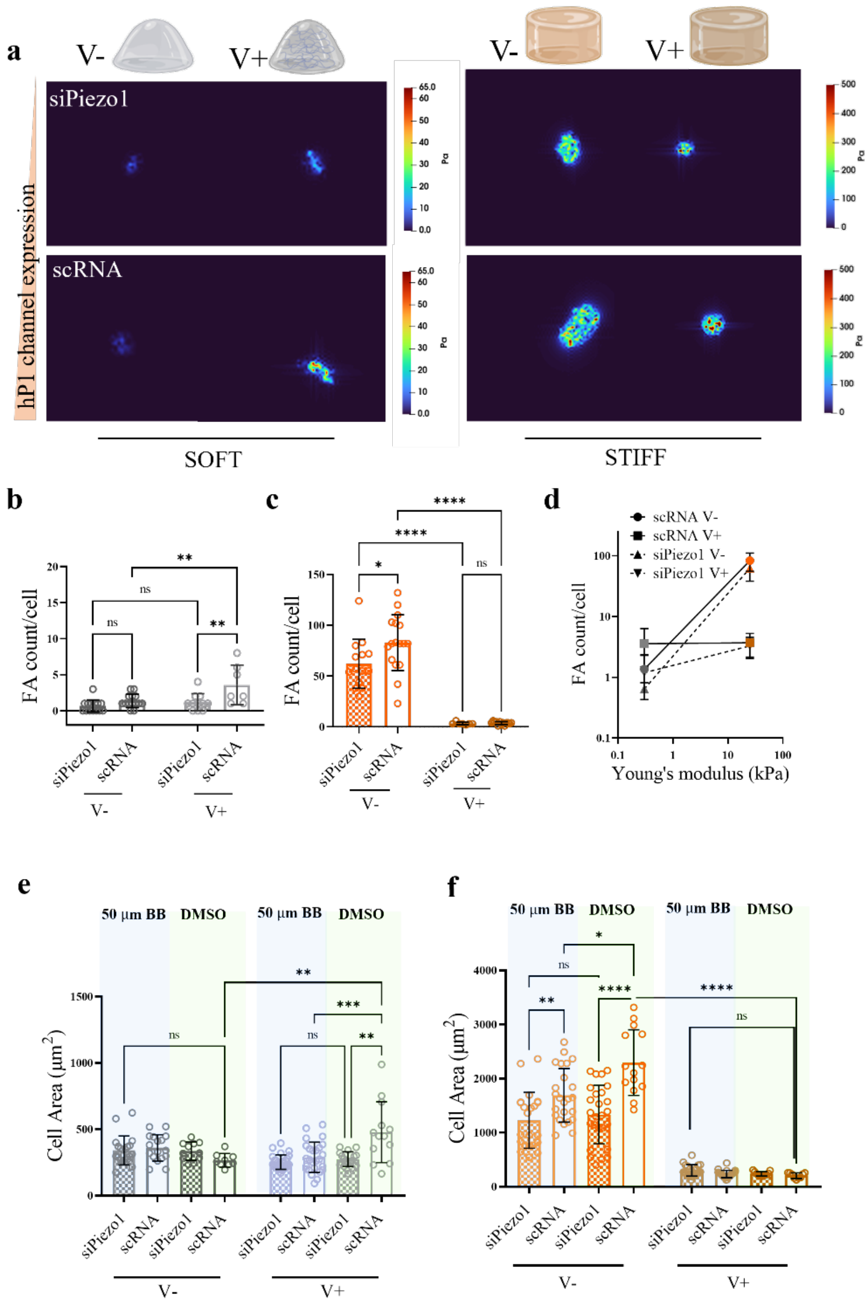


**Supplementary Fig. 3.** (**a**) Representative colour maps of traction forces applied by siPiezo1 (top) and scRNA (bottom) cells on soft and stiff hydrogel groups. (**b**) Quantified total focal adhesion per cell (FA count/cell) of the soft and stiff (**c**) hydrogel groups. Dots represent individual cell measurements, data shown as mean ± SD, n ≥ 7. (**d**) Summary of average FA count per cell ± SD plotted as a function of stiffness for all conditions. (**e**) Quantified cell area in cells with Blebbistatin (50 µM BB) treatment or without (DMSO) in the soft and (**f**) stiff group. Data shown as mean ± SD. Blue and green shaded bars represent blebbistatin and DMSO treatment conditions, respectively. P values indicating significance, ns > 0.05, * ≤ 0.05, ** ≤ 0.01, *** ≤ 0.001, **** ≤ 0.0001. N ≥ 7.

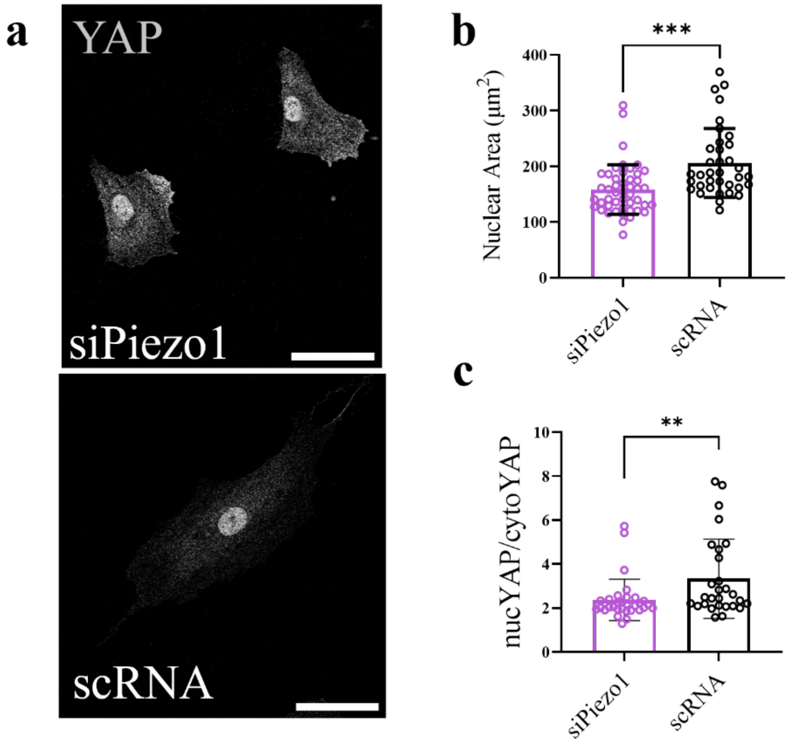


**Supplementary Fig. 4.** Representative images of YAP in (top to bottom) siPiezo1 and scRNA Y201 MSCs cultured on fibronectin coated glass coverslips for 48h. (**b**) Quantified nuclear area shown as μm2. Dots represent individual cell values and bars represent mean ± SD. (**c**) Quantified nuclear/cytoplasmic YAP ratio. Dots represent individual cell values and bars is represented as mean ± S*D,* n ≥ 36. P values indicating significance, ** ≤ 0.01, *** ≤ 0.001.

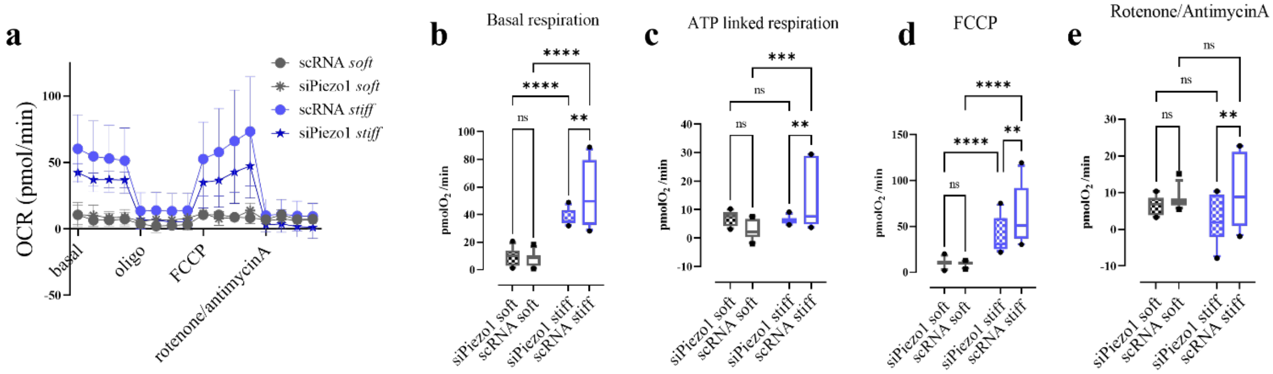


**Supplementary Fig. 5.** (**a**) Oxygen Consumption Rate (OCR) profile determined with an extracellular flux analyser of siPiezo1 and scRNA cells cultured as cell monolayers on soft ($E \sim 200$ Pa) and stiff ($E \sim 1$GPa) Matrigel coated wells. Oligomycin (oligo, 0.8 µM), FCCP (0.9 µM), rotenone (1 µM) plus antimycin A (1 µM) were used to determine the basal respiration, ATP-coupled respiration, maximal respiratory capacity, and non-mitochondrial oxygen consumption, respectively. Figure represents two independent experiments (N=2) each with n = 4 technical repeats. (**b**) Average respiration rates in the basal respiration phase of the OCR measurements. (**c**) Average respiration rate for the ATP linked respiration phase (after oligomycin addition). (**d**) Average maximal respiration capacity respiration rates after FCCP treatment. (**e**) Average non-mitochondrial respiration rates after Rotenone/Antimycin A treatment. Data shown as Box and Whiskers plot with 10-90% whiskers. A two-way ANOVA statistical test followed by a Fisher's Least Significant Difference (LSD) test was performed. P values indicating significance, ns > 0.05, ** ≤ 0.01, *** ≤ 0.001, **** ≤ 0.0001.

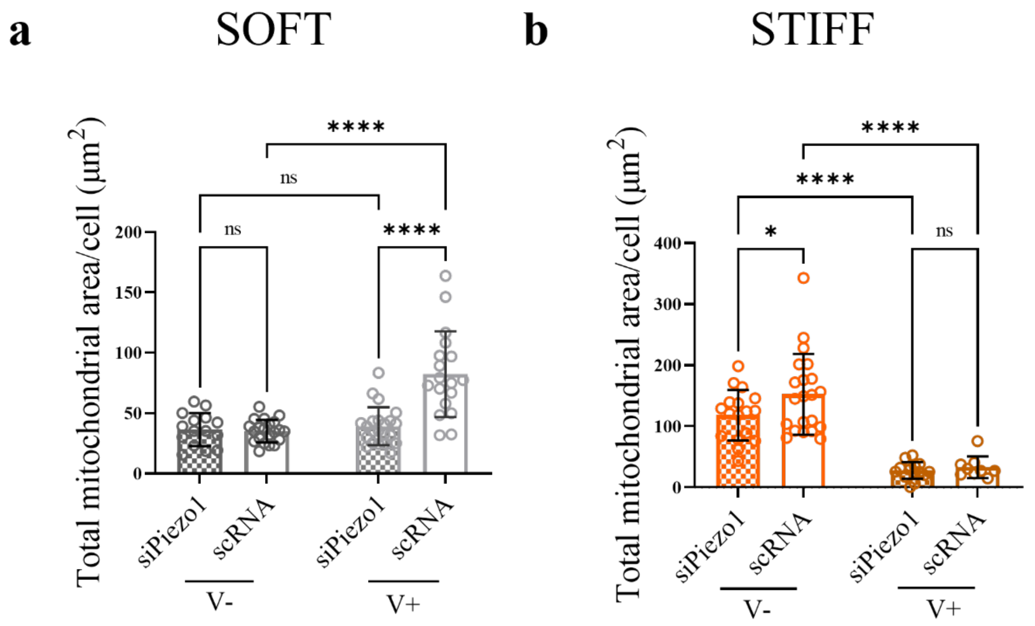


**Supplementary Fig. 6.** Quantified total mitochondrial area per cell of the soft (**a**) and stiff (**b**) hydrogel groups. All individual points represent averaged mitochondria measurements from individual cells. Data shown as mean ± SD. P values indicating significance, ns > 0.05, * ≤ 0.05, **** ≤ 0.0001, N ≥ 15.

### Supplementary Tables

**Table 1.** Hydrogel formulations.

| Reagent | Soft V- | Soft V+ | Stiff V- | Stiff V+ |
|---|---|---|---|---|
| **Aam (%)** | 3 | 3 | 15 | 35 |
| **BisAam (%)** | 0.06 | 0.06 | 0.1 | 0.0124 |
| **40% Aam (µl)** | 75 | 75 | 375 | 875 |
| **2% BisAam (µl)** | 30 | 30 | 50 | 6.2 |
| **mQ H2O (µl)** | 812.5 | / | 565 | 108.8 |
| **TEMED 1.5%/100% (µl)** | 62.5 (1.5%) | 62.5 (1.5%) | 2.5 (100%) | 2.5 (100%) |
| **APS 5%/10% (µl)** | 20 (5%) | 20 (5%) | 7.5 (10%) | 7.5 (10%) |
| **Linear Aam (µl)** | / | 812.5 | / | / |
| **Total volume (µl)** | **1000** | **1000** | **1000** | **1000** |

**Table 2**. List of antibodies and other reagents used for immunodetection.

| Reagent | Provider | Notes |
|---|---|---|
| **Alexa Fluor™ 488 Phalloidin** | Thermo (A12379) | 1:250 |
| **Anti-YAP Antibody (63.7)** | Santa Cruz (sc-101199) | 1:100 |
| **Anti-Fibronectin antibody** | Sigma (F3648) | 1:200 |
| **Anti-Vinculin antibody** | Sigma (V9264) | 1:400 |
| **Anti-TOMM20** | Abcam (ab186735) | 1:200 |
| **Anti-Piezo1** | Sigma (AMAB91589) | 1:100 |
| **Cy™3 AffiniPure Rabbit Anti-Mouse IgG (H+L)** | Jackson ImmunoResearch (315-165-003) | 1:200 |
| **Donkey anti-Rabbit IgG (H+L) Alexa Fluor™ 488** | Invitrogen (A-21206) | 1:250 |
| **CellTag™ 700 Stain for In-Cell Western™ Assays** | Li-Cor (926-41090) | 1:500 |
| **IRDye® 800CW Goat anti-Mouse IgG Secondary Antibody** | Li-Cor (926-32210) | 1:500 |
